# Inhibition of Stemness Pathways during Acute SIV Limits Infection of Central Memory CD4+ T Cells and Alters Viral Reservoir Activity in Macaques

**DOI:** 10.64898/2026.07.02.734630

**Authors:** Riri Rizkianty Hamid, Inna Ruiz-Salinas, Nils Schoof, Alora Colvin, Sijia Tao, Selwyn J. Hurwitz, Alice Lin, Jordan Goldy, Gregory K Tharp, Steven E Bosinger, Raymond F. Schinazi, Guido Silvestri, Ann Chahroudi, Maud Mavigner

**Affiliations:** Department of Pediatrics, Emory University School of Medicine, Atlanta, Georgia, USA; Center for ViroScience and Cure, Laboratory of Biochemical Pharmacology, Department of Pediatrics, Emory University School of Medicine and Children’s Healthcare of Atlanta, Atlanta, Georgia, USA; Emory National Primate Research Center, Emory University, Atlanta, Georgia, USA; Center for Childhood Infections and Vaccines of Children’s Healthcare of Atlanta and Emory University, Atlanta, Georgia, USA

## Abstract

The establishment of a reservoir of latently infected CD4+ T cells that persist on antiretroviral therapy (ART) through proliferation represents the main barrier to HIV cure. Here, we evaluated in macaques a therapeutic approach targeting Wnt and Notch pathways that regulate T cell proliferation and differentiation during acute SIV infection, when the viral reservoir is seeded. The combination of PRI-724 and LY3039478 led to reprogramming of central memory CD4+ T cells away from quiescence and stemness toward a metabolically active effector profile resulting in reduced infection of central memory CD4+ T cells. Following sustained ART, levels of SIV RNA in CD4+ T cells were higher in the PRI-724 + LY3039478-treated group compared to controls, although SIV DNA was similar. These findings suggest that stemness pathway inhibition promotes memory T cell differentiation leading to a more transcriptionally active reservoir and has potential to synergize with “shock-and-kill” approaches to reduce HIV persistence.

## Introduction

While antiretroviral therapy (ART) has markedly improved the prognosis for people with HIV, lifelong treatment remains necessary due to the persistence of a reservoir of latently infected CD4+ T cells generated at a very early stage of the infection^1,2^ and maintained over time through cellular proliferation^3–6^. HIV establishes latency in all maturational subsets of CD4+ T cells, including naïve, stem cell memory (SCM), central memory (CM), transitional memory (TM), and effector memory T cells (EM) which each contribute differentially to HIV persistence^7–10^. While cells displaying a differentiated phenotype such as EM carry most clonally expanded HIV proviruses persisting on ART, these cellular clones may wane overtime^11,12^. Additionally, these cells appear to be the progeny of less differentiated SCM and CM cells that constitute a stable core of the HIV reservoir due to their stem cell-like properties of intrinsic high survival and self-renewal abilities, long half-lives, and multipotency^10,13–21^.

The fate of these long-lived immature T cells to self-renew or differentiate is regulated by a network of intricate biological pathways^22–25^, all representing potential therapeutic targets to disrupt HIV persistence. Modeling studies of HIV dynamics indeed suggest that therapies inhibiting T cell proliferation and/or promoting T cell differentiation could substantially reduce HIV reservoir size^23,26,27^. The Wnt/β-catenin signaling plays a well-established role in the generation and maintenance of memory T cells^10,14,28,29^. Activation of this pathway by ligand-receptor interaction triggers the release, stabilization, and translocation of β-catenin to the nucleus where it interacts with members of the T cell factor (TCF) family and transcriptional co-activators to activate target gene transcription. The co-activator p300 and its homologue CREB-binding protein (CBP) act as bimodal regulators promoting differentiation or self-renewal transcriptional programs respectively^30^. We previously demonstrated that pharmacological modulation of the Wnt/β-catenin signaling pathway results in inhibition of proliferation and induction of differentiation of long-lived memory CD4+ T cells in ART-treated rhesus macaques (RMs)^31^. Specifically, we found that PRI-724, a small molecule that blocks the interaction between CBP and β-catenin, reduced the proliferation of SCM and CM CD4+ T cells and promoted a transcriptome enriched in differentiation genes. However, PRI-724 treatment alone during ART-suppressed SIV infection did not significantly impact viral reservoir size or distribution, suggesting that a combination approach targeting multiple signaling pathways may be needed to achieve these outcomes.

The Notch pathway, similar to the Wnt pathway, is a key regulator of stem cell and T cell quiescence, self-renewal, and differentiation^24,32–34^. Notch receptor-ligand interaction triggers a two-step proteolytic cleavage of the receptor mediated first by a disintegrin and metalloproteinase (ADAM) enzyme, and second by a γ-secretase. The resulting free Notch intracellular domain translocates into the nucleus where it interacts with nuclear factors including CBP and p300 to regulate target gene expression^34,35^. Although Wnt and Notch are separate pathways, these two pathways are complimentary in regulating cell self-renewal, quiescence, and cell fate^25,35–37^. Inhibitors of the γ-secretase have been developed for oncologic indications and studies have shown that they block proliferation and induce apoptosis of cancer stem cells thus inducing tumor regression^38,39^. In addition, the γ-secretase inhibitor LY3039478 has been found to be safe in several clinical trials^40–43^.

In the current study we assessed the impact of the combined inhibition of Wnt and Notch signaling on CD4+ T cell dynamics and virus distribution during acute SIV infection as well as reservoir stability during ART in RMs. We found that treatment with the CBP/β-catenin inhibitor PRI-724 and the γ-secretase inhibitor LY3039478 induced profound transcriptional changes in CM CD4+ T cells indicative of a shift toward effector differentiation accompanied by a transient reduction of their contributions to both the CD4+ T cell and infected cell pools. Although the reservoir size in CD4+ T cells was similar in Wnt- and Notch-inhibited and control RMs following ART suppression, the reservoir was more transcriptionally active several months after completion of the experimental treatments. These findings suggest that inhibiting the Wnt and Notch stemness pathways during acute infection leads to a less stable reservoir, providing a potential window of opportunity for latency reversal and immune clearance strategies.

## Results

### Notch inhibitor dose-finding study

To target Wnt and Notch pathways we chose to combine two drugs investigated in several oncological clinical trials^41,43–45^, the CBP/β-catenin inhibitor PRI-724 used in our previous study in ART-suppressed SIV-infected RMs^31^, and the γ-secretase inhibitor LY3039478. In order to select the dose of LY3039478 to be used in vivo in combination with PRI-724, we performed a dose-escalation study in two uninfected rhesus macaques (RMs) consisting of two 8-week treatment cycles of PRI-724 at the previously defined dose of 20 mg/kg daily^31^ and LY3039478 oral administration at escalating doses of 1.5 and 2.5 mg/kg three times a week (Supplementary Fig. 1A). Plasma levels of C-82, the active metabolite of PRI-724^44,46,47^, and LY3039478 were measured by liquid chromatography–mass spectrometry (LC-MS/MS) over 24 hours following drug administration. A consistent pharmacokinetic profile for C-82 between animals and treatment cycles was observed. Plasma exposure of LY3039478 at the higher dose of 2.5 mg/kg appeared to increase more than dose-proportionally, reaching pharmacokinetic levels comparable to those reported in clinical trials^41,43,48^ (Supplementary Fig. 1B, C).

In addition to the cleavage of the Notch receptor, γ-secretase has various other targets including the amyloid-β precursor protein (APP). As APP cleavage results in amyloid-β accumulation, measuring its plasma levels can serve as a pharmacodynamic marker of LY3039478 activity^49^. Administration of LY3039478 at the increased dose of 2.5 mg/kg resulted in a transient reduction in plasma levels of amyloid-β, both after the first dose and after repeated dosing (Supplementary Fig.1D), thus confirming the biological activity of the Notch inhibitor in our RM model. Furthermore, the experimental treatments were well tolerated in healthy RMs with no significant impact on complete blood counts or serum chemistries and no adverse events observed (Supplementary Fig. 2). Given the acceptable safety, pharmacokinetic and pharmacodynamic profiles of the PRI-724 (20 mg/kg) and LY3039478 (2.5 mg/kg) combination, this regimen was selected for administration to SIV-infected RMs.

**Fig. 1.**
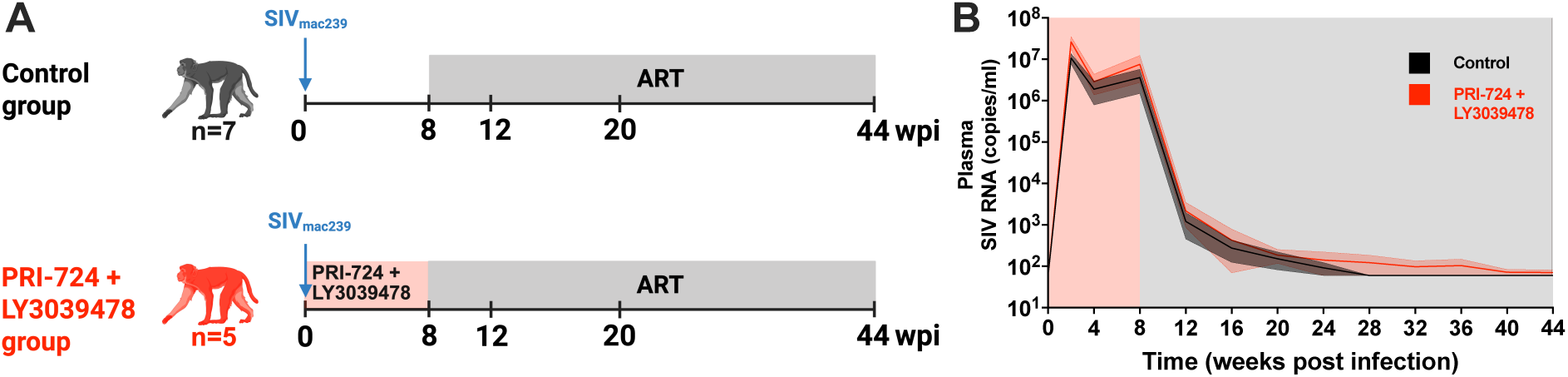
Study design and plasma viral load kinetics. (**A**) Experimental design. Five RMs were infected intravenously with 10^5^ TCID_50_ (50% tissue culture infective dose) of SIVmac239 before receiving the combined treatment PRI-724 (20 mg/kg, subcutaneously, daily) + LY3039478 (2.5 mg/kg, orally, three times per week) for 8 weeks. ART was initiated 8 weeks post infection (wpi) and maintained for the duration of the study. Seven RMs were treated with ART only and served as controls. (**B**) Longitudinal assessment of plasma SIV gag RNA levels in PRI-724 + LY3039478-treated RMs (red) and controls (black). The red shaded area represents the PRI-724 + LY3039478 combined treatment period. The grey shaded area represents the ART treatment period. Data is represented as mean ± standard error of the mean.

### Experimental design of the efficacy study

To evaluate whether pharmacological modulation of stemness signaling pathways destabilizes HIV/SIV reservoir establishment, we treated 5 RMs with PRI-724 + LY3039478 immediately following SIV infection for 8 weeks until ART initiation (Fig. 1A). Seven RMs were infected with SIV and treated with ART only as control. All 12 RMs were males of Indian-origin and negative for the Mamu-B*08 and Mamu-B*17 major histocompatibility complex (MHC) class I alleles associated with an increased frequency of spontaneous control of SIV replication^50,51^. SIV challenge consisted of a single intravenous (i.v.) administration of 1x10^5^ 50% tissue culture infective doses (TCID_50_) of SIVmac239. ART, consisting of tenofovir disoproxil fumarate (TDF), emtricitabine (FTC) and dolutegravir (DTG) was administered daily by subcutaneous (s.c.) injection from 8 weeks post infection (wpi) to the end of the study.

Plasma viral loads were similar between groups with peaks ranging from 3.2x10^6^ to 4.8x10^7^ copies of SIV gag RNA per ml of plasma (Fig. 1B and Table 1). Clinical and laboratory parameters were assessed longitudinally to determine the safety and tolerability of PRI-724 and LY3039478 administration during acute SIV infection. There were no adverse events and only transient elevations in liver enzymes during the period of combined treatment with PRI-724 + LY3039478 (Supplementary Fig. 3).

**Table 1.** Characteristics of study groups. Statistical significance was determined with a two-sided Mann-Whitney U-test.

| Group | ID | Sex | Age at infection (years) | Peak PVL (copies/ml) |
| --- | --- | --- | --- | --- |
| Control | RFg16 | Male | 3.5 | $2.9 \times 10^7$ |
| | RAr16 | Male | 1.6 | $4.8 \times 10^7$ |
| | RUs16 | Male | 2.6 | $4.1 \times 10^7$ |
| | REi16 | Male | 2.7 | $1.1 \times 10^7$ |
| | RQf16 | Male | 3.5 | $3.2 \times 10^6$ |
| | RKn16 | Male | 2.6 | $1.9 \times 10^7$ |
| | RAj16 | Male | 2.7 | $2.4 \times 10^7$ |
| | | Mean | 2.7 | $1.9 \times 10^7$ |
| PRI-724 +<br>LY3039578 | REg18 | Male | 4.7 | $1.1 \times 10^7$ |
| | RYg18 | Male | 4.5 | $3.2 \times 10^6$ |
| | RHn19 | Male | 3.0 | $4.8 \times 10^7$ |
| | RPj19 | Male | 3.0 | $2.9 \times 10^7$ |
| | RYp19 | Male | 2.9 | $4.1 \times 10^7$ |
| | | Mean | 3.6 | $3.2 \times 10^7$ |
|  |  | Significance | n.s | n.s |

### Inhibition of Wnt and Notch pathways reduces the contribution of central memory cells to the peripheral CD4+ T cell compartment

During the first 8 weeks of SIV infection (pre-ART), both groups of animals displayed increasing relative frequencies of naïve CD4+ T cells due to the preferential loss of memory CD4+ T cells. Significantly lower frequencies of central memory (CM) CD4+ T cells were observed at 8 wpi in RMs treated with PRI-724 + LY3039478 compared to controls (P=0.0051, Fig. 2A). A more pronounced reduction from pre-infection of CM CD4+ T cells in the PRI-724 + LY3039478-treated group was found compared to controls (P=0.0101, Fig. 2B). The absolute counts of CM CD4+ T cells were also lower in the PRI-724 + LY3039478-treated group at 8 wpi (P=0.0101, Supplementary Fig. 4A) with a more drastic decline following infection as compared to controls (P=0.0303, Supplementary Fig. 4B). Despite a greater recovery in CM CD4+ T cell frequencies and absolute counts following ART initiation in the PRI-724 + LY3039478-treated group versus control group (P=0.0479, Fig. 2B and P=0.0177, Supplementary Fig. 4B), the frequency of CM CD4+ T cells remained lower in the PRI-724 + LY3039478-treated group at 12 wpi (P=0.0177, Fig. 2A). Interestingly, after several months of ART, subsets with intermediate/late differentiation status showed higher levels in the PRI-724 + LY3039478-treated group as compared to controls. This difference reached statistical significance for the frequency of TM CD4+ T cells (P=0.0177, Fig. 2A) and trended towards significance for EM CD4+ T cells. Altogether, these results suggest an impact of stemness pathway inhibition on CD4+ T cell dynamics characterized by a decreased contribution of the CM subset to the circulating CD4+ T cell pool.

**Fig. 2.**
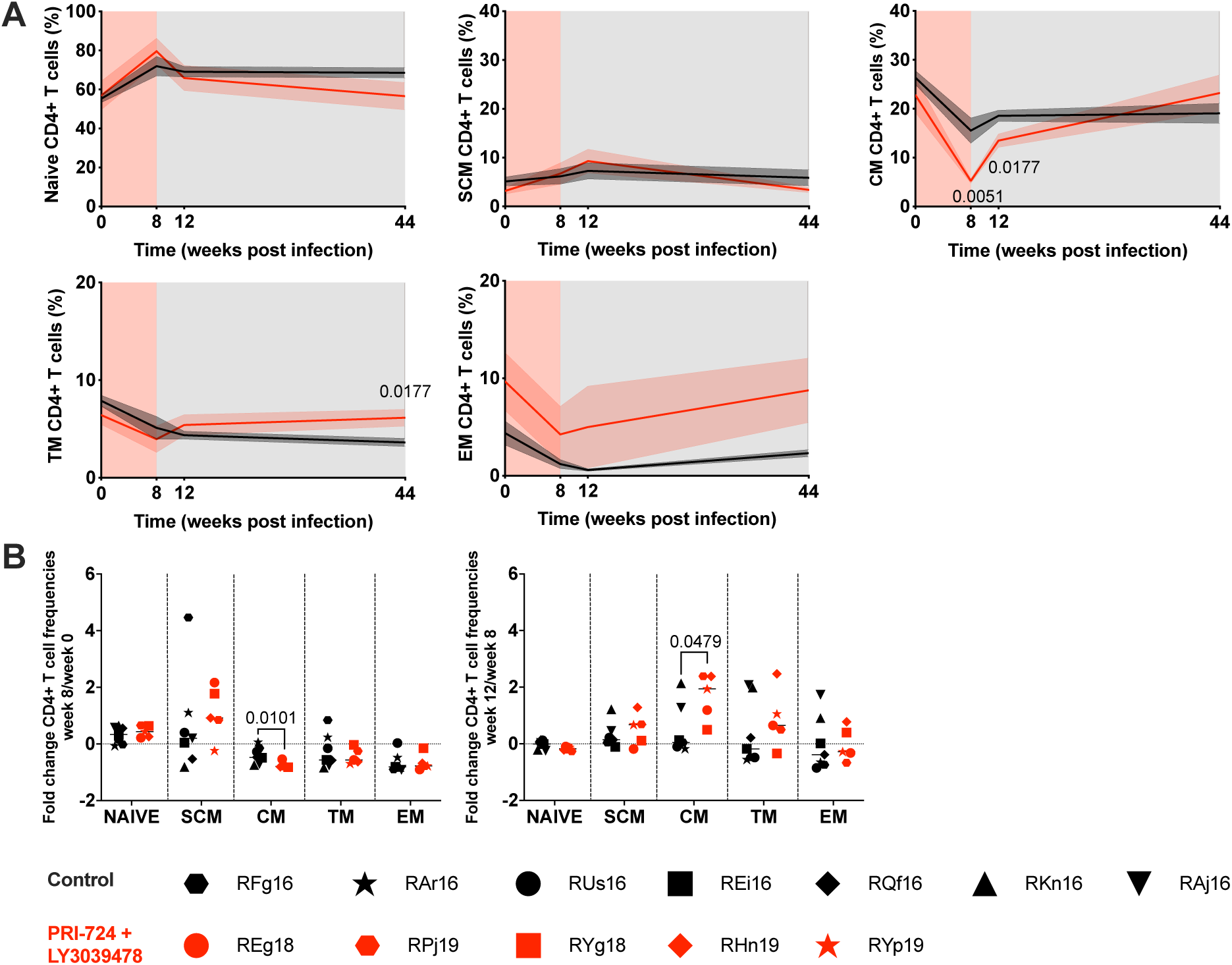
Impact of PRI-724 + LY3039478 combined treatment on CD4+ T cell subset frequency. (**A**) Longitudinal assessment of the contribution of CD4+ T cell subsets to the total peripheral CD4+ T cell pool in PRI-724 + LY303947-treated RMs (red) versus controls (black). The red shaded area represents the PRI-724 + LY3039478 combined treatment period. The grey shaded area represents the ART treatment period. Data is represented as mean ± standard error of the mean. (**B**) Fold change of CD4+ T cell frequencies between 0 and 8, and between 8 and 12 week post infection. Horizontal lines represent the median. A two-sided Mann-Whitney U-test was used to compare values between groups.

### Inhibition of Wnt and Notch pathways promotes an activated effector-like profile in CM CD4+ T cells

To gain insight into the mechanisms supporting the PRI-724 + LY3039478-induced reduction in CM CD4+ T cells, we performed RNA-seq analyses of purified CM cells sorted prior to and immediately following the stemness inhibitor treatment period, and at equivalent timepoints for the controls (gating strategy shown in Supplementary Fig. 5). In the control group, we observed 169 differentially expressed genes (DEGs) between 0 and 8 wpi (Figure 3A), which is not surprising given the major perturbation in the T cell compartment during the acute phase of SIV infection. In contrast, in the PRI-724 + LY3039478-treated group, we observed a more than 10-fold increase in DEGs, finding 1,790 transcripts differentially expressed between 0 and 8 wpi (Fig. 3A). We formally tested for genes significantly higher or lower after PRI-724 + LY3039478 treatment compared to those changing in the conrol group due to acute SIV infection alone using an interaction analysis, and found that 267 genes were significantly differentially regulated in CM CD4+ T cells from PRI-724 + LY3039478-treated RMs as compared to their transcriptional changes in controls (Fig. 3A, left box). Amongst the upregulated genes in PRI-724 + LY3039478-treated animals versus controls, were the transcriptional regulators NRIP1 and TRAF6 (Fig. 3B). TRAF6 encodes an adapter protein known to be crucial for T cell activation and effector functions^52,53^. The protein encoded by NRIP1 inhibits the Wnt pathway and has been reported to limit the pool of stem-cell-like T cells and restrict memory formation^54^.

**Fig. 3.**
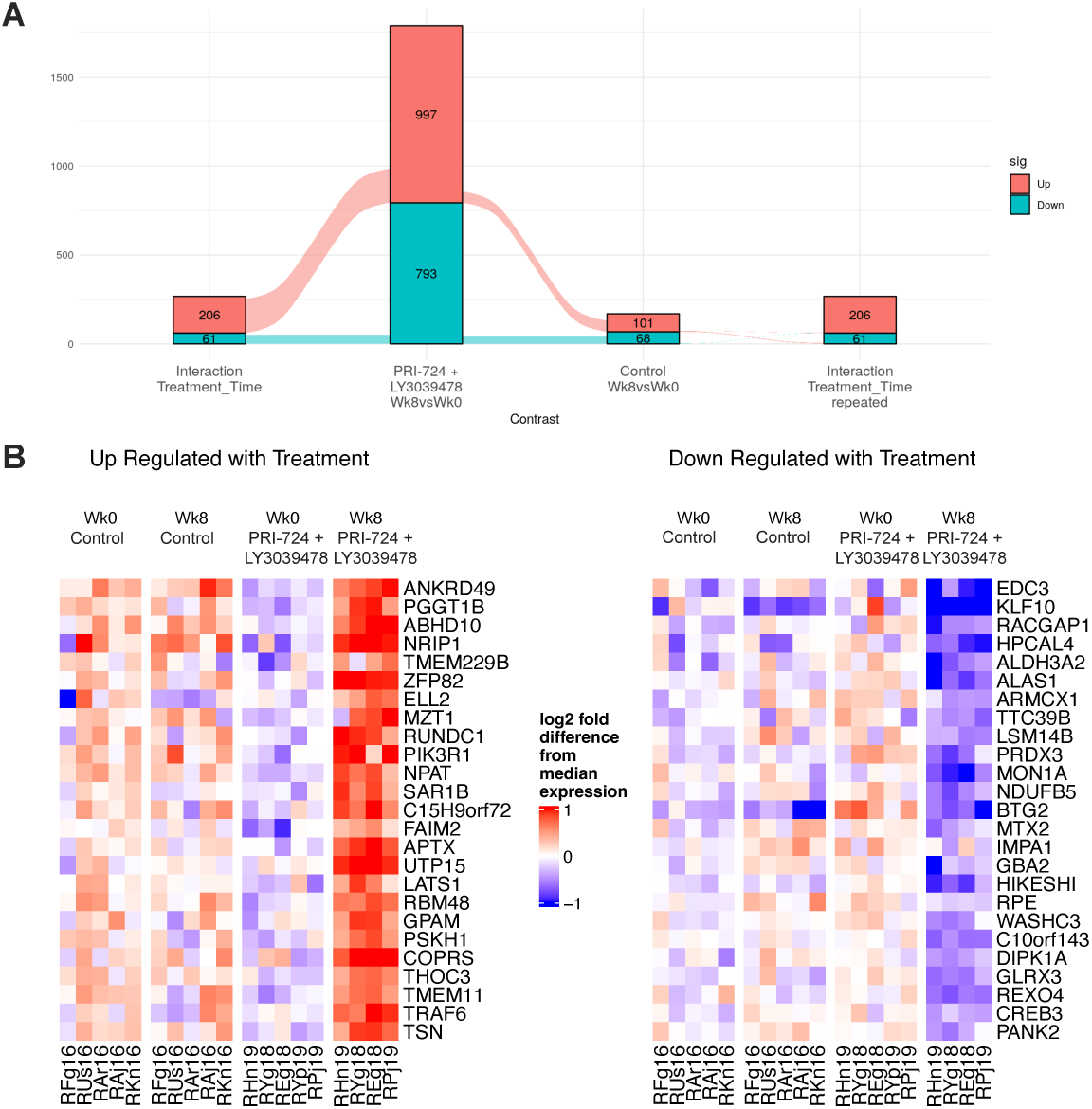
Impact of PRI-724 + LY3039478 combined treatment on CM CD4+ T cell whole transcriptomic profile. Circulating CM CD4+ T cells isolated by flow sorting from PRI-724 + LY3039478-treated and control RMs at 0 and 8 weeks post infection were analyzed by bulk RNA-Seq. (**A**) Alluvial plot comparing the number of differentially expressed genes (DEG) in control and experimental groups over the treatment period. Ribbons indicate number of genes in common in the comparison. The left-end and right-end boxes show the DEGs in the interaction analysis contrasting the PRI-724 + LY3039478 treatment effect to the genes changing over time in the control animals; the box is repeated on each end to show the overlap in genes with the treatment and control analyses. The two center boxes indicate the direct constrast of week 8 vs week 0 for the PRI-724 + LY3039478-treated and control groups. (**B**) Heatmap of the leading-edge genes differentially regulated between week 0 and 8 post infection in CM CD4+ T cells from PRI-724 + LY3039478-treated RMs as compared to controls (at least 2-fold differential expression; adjusted P value of 0.05).

We next performed gene set enrichment analysis (GSEA) using gene sets from the reactome and hallmark databases. Over the PRI-724 + LY3039478 treatment period, we observed an upregulation of genes associated with the cell cycle, notably components of the proteasome, in PRI-724 + LY3039478-treated RMs as compared to controls (Fig. 4A). Furthermore, CM CD4+ T cells from animals in the PRI-724 + LY3039478-treated group showed increased glycolysis and lipid metabolism and decreased oxidative phosphorylation (OXPHOS) and amino acid degradation compared to controls (Fig. 4B). This metabolic profile is indicative of T cell activation and compatible with effector T cell differentiation^55–57^. Interestingly, it has been shown that HIV downregulates glycolysis to enter a latent state, and that restoration of the glycolytic activity is required for HIV reactivation^58^.

**Fig 4.**
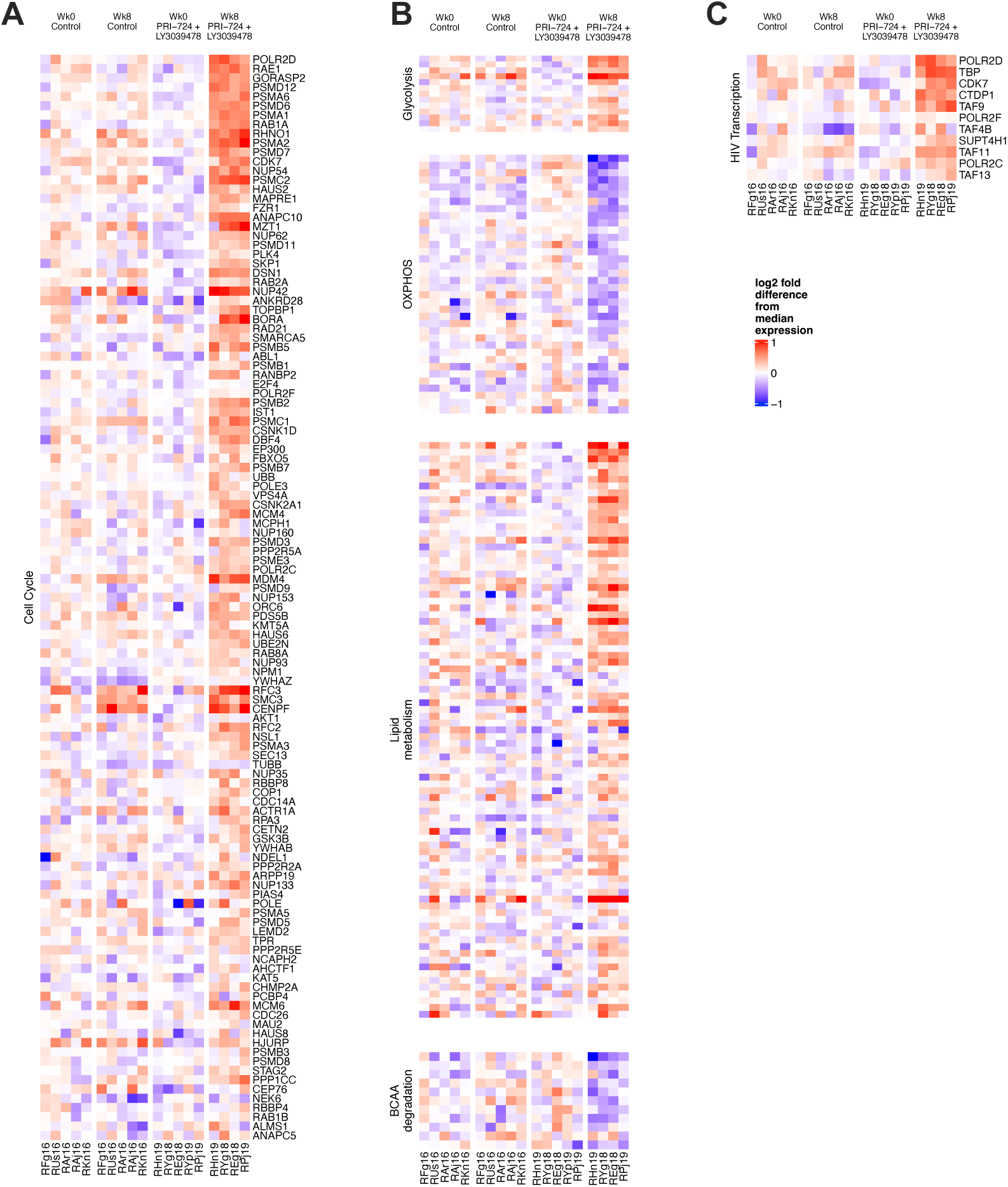
Impact of PRI-724 + LY3039478 combined treatment on CM CD4+ T cell gene set expression. A-C. Heatmaps showing Z score gene expression from (**A**) cell cycle, (**B**) cell metabolism, and (**C**) HIV/SIV transcription gene sets differentially regulated between week 0 and 8 post infection in sorted circulating CM CD4+ T cells from PRI-724 + LY3039478-treated RMs as compared to controls (at least 2-fold differential expression; adjusted P value of 0.05). OXPHOS: oxidative phosphorylation; BCAA: branched-chain amino acids.

The transcriptomic analyses further revealed an upregulation of genes associated with HIV transcription including genes coding for RNA polymerase and other components of the transcription machinery (Fig. 4C). This observation is consistent with the upregulation of TRAF6 that activates HIV transcription through Tat-induced activation of NF-κB^59^. Consistent with their transcriptomic profile, CM CD4+ T cells from the PRI-724 + LY3039478-treated RMs also expressed higher levels of the activation marker HLA-DR at 8 wpi as compared to controls (P=0.0025, Supplementary Fig. 6). Overall, these results show that treatment with Wnt and Notch inhibitors induced a shift of the CM CD4+ T cell profile toward a less quiescent, effector-like, metabolically active state that may be less favorable to SIV latency.

### Inhibition of Wnt and Notch pathways alters SIV reservoir distribution within CD4+ T cell maturational subsets

Given the impact of PRI-724 + LY3039478 on CM frequencies and gene expression, we next wanted to assess whether stemness pathway inhibition during acute SIV modulated infection of CD4+ T cells. Subsets of naïve, SCM, CM, TM and EM CD4+ T cells were sorted by FACS from the peripheral blood of all 12 RMs at 8, 12, 20 and 44 wpi (gating strategy shown in Supplementary Fig. 5). Levels of cell-associated unintegrated SIV 2-LTR and gag DNA were quantified by multiplex quantitative PCR with the cell input determined by quantification of the RPP30 host gene. In the absence of ART, at 8 wpi, high levels of unintegrated SIV 2-LTR and gag DNA were detected across all subsets of CD4+ T cells with no significant differences between the PRI-724 + LY3039478 and control groups (Fig. 5A and 5B). During this initial phase of SIV infection pre-ART, viral DNA levels were positively correlated within subsets and with the plasma viral load with a clear exception for 2-LTR levels in TM CD4+ T cells in the PRI-724 + LY3039478-treated group (Supplementary Fig. 7).

**Fig 5.**
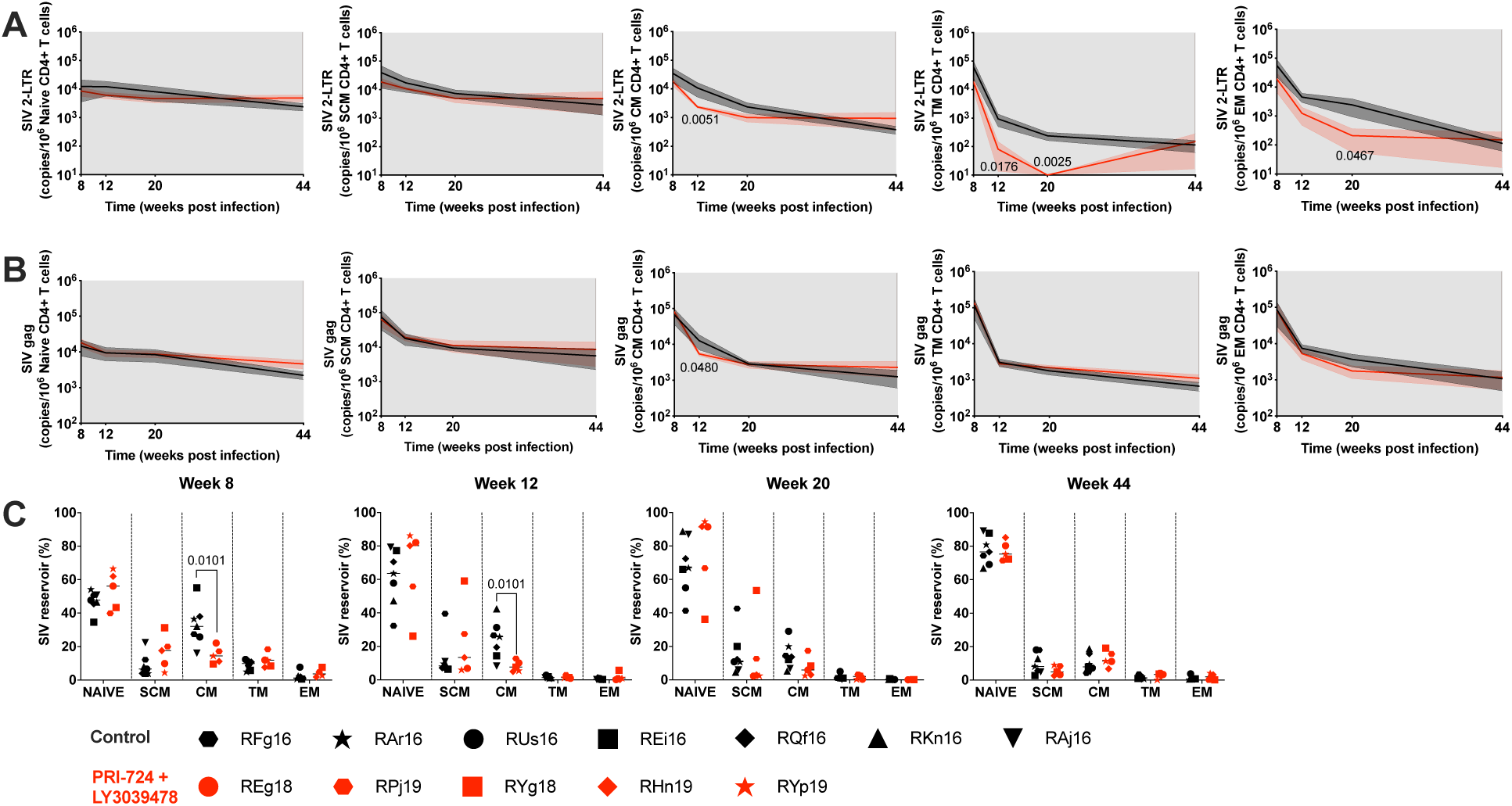
Impact of PRI-724 + LY3039478 combined treatment on SIV DNA distribution in CD4+ T cell subsets. A-C. Peripheral CD4+ T cell subsets at 8, 12, 20, and 44 weeks post infection in PRI-724 + LY3039478-treated RMs (red) versus controls (black). Longitudinal assessment of SIV (**A**) 2-LTR levels and (**B**) gag total DNA levels. The grey shaded area represents the ART treatment period. Data is represented as mean ± standard error of the mean. (**C**) Relative contribution of CD4+ T cells subsets to the SIV reservoir. Horizontal bars represent the median. A two-sided Mann Whitney U-test was used to compare values between groups.

As expected, levels of SIV 2-LTR and gag DNA decreased in all subsets of CD4+ T cells following ART initiation with a sharper decline during the first month of ART (8-12 wpi) observed in more differentiated memory cells that have high turnover rates and are more prone to cell death^27^ (Supplementary Fig. 8). RMs treated with stemness inhibitors exhibited lower levels of 2-LTR DNA in the CM (P=0.0051, 12 wpi), TM (P=0.0176, 12 wpi and P=0.0025, 20 wpi) and EM CD4+ T cells (P=0.0467, 20 wpi) as compared to controls. These results suggest that treatment with PRI-724 + LY3039478 may predispose these cells to divide or die (Fig. 5A), as 2-LTR circles decrease during ART due to the death of the cells that carry them but also through dilution during cell division^60^. Interestingly, ART initiation was followed by a greater decrease in SIV gag DNA in CM CD4+ T cells in the PRI-724 + LY3039478-treated group compared to controls resulting in significantly lower levels of SIV gag DNA at 12 wpi (P=0.0480, Fig. 5B).

When considering the contribution of each memory subset to the total SIV-infected peripheral CD4+ T cell population, a reduced contribution of CM to the total SIV gag DNA content in CD4+ T cells was found in the PRI-724 + LY3039478-treated RMs as compared to controls at both 8 and 12 wpi (P=0.0101 for both, Fig. 5C). Altogether, these results suggest an impact of stemness inhibitor treatment on SIV DNA seeding by transiently reducing the relative contribution of the CM CD4+ T cells to the peripheral CD4+ T cell SIV reservoir.

### Inhibition of Wnt and Notch pathways promotes a more transcriptionally active HIV reservoir but does not reduce its size

Having demonstrated an impact of stemness inhibitor treatment on the early distribution of SIV DNA among memory CD4+ T cell subsets, we sought to compare viral persistence in CD4+ T cells between groups after several months of viral suppression on ART. Significant differences were not observed in the levels of SIV gag DNA in peripheral CD4+ T cells in the PRI-724 + LY3039478-treated versus control group at 36 and 44 wpi (Fig. 6A). We noted that for the control RMs the level of SIV DNA persisting after several months of ART was positively correlated with the pre-ART plasma viral load (R=0.7500, P=0.0663 at 36 wpi; R=0.8829, P=0.0151 at 44 wpi, Fig. 6B). This correlation was not found in the PRI-724 + LY3039478-treated group reinforcing the previous observation of disrupted reservoir seeding by the experimental treatment even if long term reductions in SIV DNA persistence were not seen. At both 36 and 44 wpi, higher levels of SIV gag RNA in CD4+ T cells from RMs treated with PRI-724 + LY3039478 versus controls were found (P=0.0303 and P=0.0480, Fig. 6C). This result indicates that PRI-724 + LY3039478 treatment during acute SIV infection leads to a more transcriptionally active and potentially less stable viral reservoir.

**Fig. 6.**
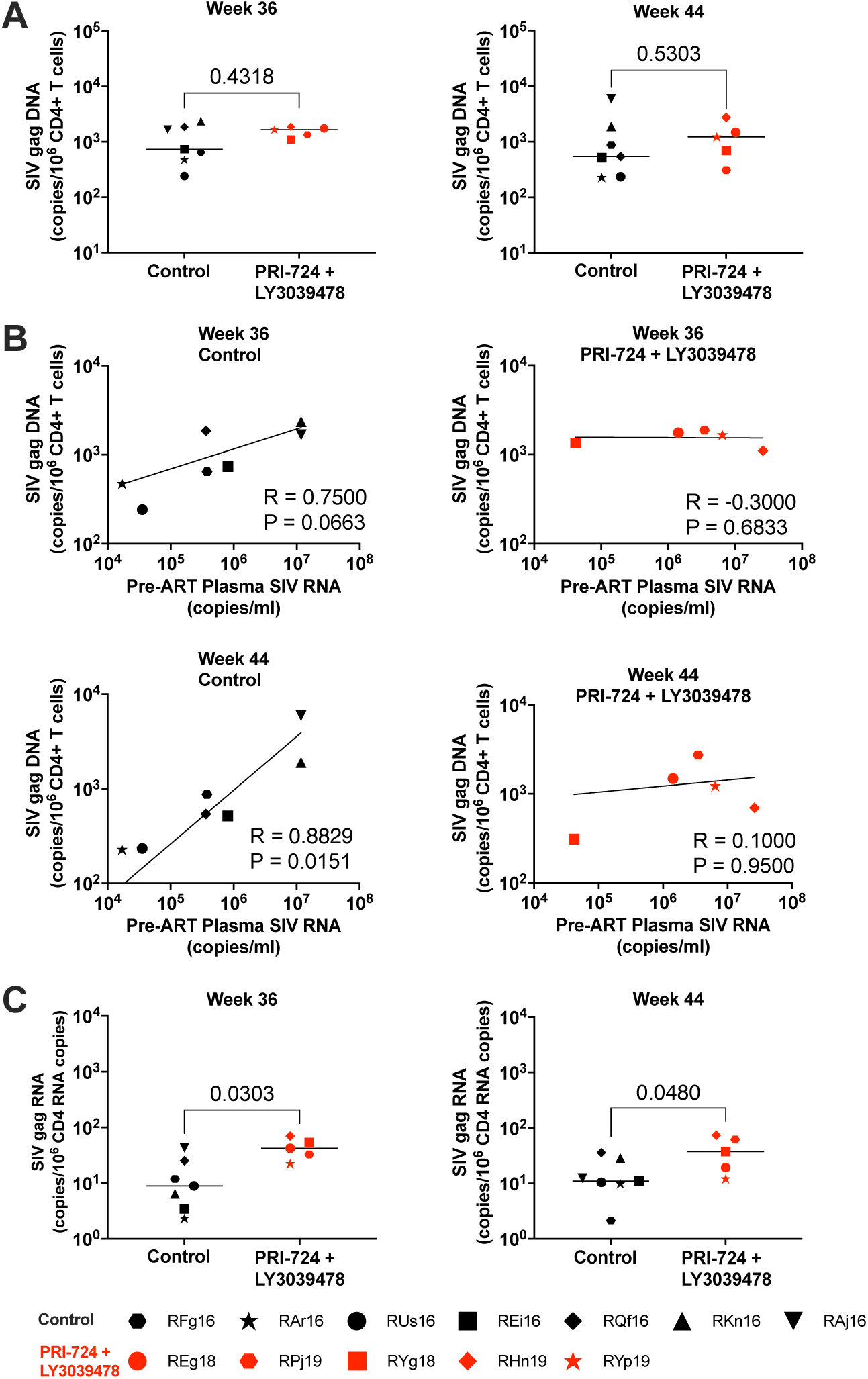
Impact of PRI-724 + LY3039478 combined treatment on SIV persistence in total CD4+ T cells. (**A**) SIV gag DNA level in total circulating CD4+ T cells after extended ART and (**B**) its association with pre-ART plasma viral load and (**C**) SIV gag RNA level in total circulating CD4+ T cells after extended ART. Horizontal lines represent the median. Two-sided Mann-Whitney U-tests and Spearman correlation tests were used.

## Discussion

The main barrier to HIV cure is the establishment of a latent infection in a heterogenous population of CD4+ T cells that persists during ART. The maintenance of this latent reservoir over time is defined by biological processes regulating the lifespan, proliferation, and differentiation of CD4+ T cells^61–63^. Here we targeted latent reservoir establishment with a pharmacological approach to inhibit CD4+ T cell self-renewal and promote differentiation using Wnt and Notch signaling pathway inhibitors. Given their pleiotropic functions, modulating stemness pathways in vivo requires caution. Previously, we demonstrated an acceptable safety profile of the Wnt inhibitor PRI-724 in ART-suppressed SIV-infected RMs^31^. Here we additionally report an absence of adverse effects associated with the combined administration of the Notch inhibitor LY3039478 with PRI-724 during acute SIV infection of RMs. We found that combined Wnt and Notch inhibition during acute SIV infection induced a shift in CM CD4+ T cells toward a metabolically active effector-like profile, reduced their infection, and increased transcriptional activity of the CD4+ T cell reservoir following sustained ART.

Our approach targeting stemness pathways during acute SIV infection had the greatest effect on CM CD4+ T cells, with a transient reduction of their frequencies and absolute counts immediately following experimental treatment and maintained during the first few weeks of ART initiation. Our previous study using the Wnt inhibitor PRI-724 alone in ART-suppressed SIV-infected RMs also demonstrated significant treatment-related modulation of the transcriptomic profile in CM and SCM CD4+ T cells specifically, but not in naive or EM CD4+ T cell subsets^31^. A specific impact on less mature long-lived cells was expected as Wnt and Notch pathways are more active in cells presenting enhanced self-renewing abilities^14,64,65^. The more pronounced decline in CM CD4+ T cells we observed in the experimental group may be the result of enhanced cell death induced by the stemness pathway inhibition and/or from their increased differentiation into TM CD4+ T cells. Although not mutually exclusive, the latter option is supported by the observed increased frequency of TM CD4+ T cells and to a lesser extent EM CD4+ T cells in the experimental group as compared to controls. Increased CD4+ T cell differentiation is also supported by our transcriptomic and phenotyping analyses showing clear alterations of the CM CD4+ T cell profile in terms of activation and metabolism. CD4+ T cells with stem-like properties such as CM CD4+ T cells balance self-renewal with effector differentiation to maintain immune memory while replenishing the short-lived effector cell compartment. This dynamic process is concomitant with drastic changes in cellular metabolism. When transitioning from a quiescent, stem-like state to an activated, effector state, T cells underdo a metabolic switch from OXPHOS to glycolysis to satisfy the high energy demands of effector functions^66,67^. This metabolic switch was observed in CM CD4+ T cells following experimental treatment of the RMs with Wnt and Notch inhibitors.

The consequences of the stemness inhibitor-induced changes affecting CM CD4+ T cells on SIV reservoir seeding and persistence were evaluated by a longitudinal assessment of total and unintegrated viral levels in each maturational subset. The decline of CM CD4+ T cells in the PRI-724 + LY3039478-treated group was accompanied by lower total SIV DNA levels and a reduced contribution of CM to the CD4+ T cell reservoir. Preferential sparing of CM CD4+ T cells from SIV/HIV infection has been associated with better viral control, reduced immunopathology and decreased risk of disease progression^68,69^. This is notably a key characteristic of natural hosts of SIV infection such as Sooty Mangabeys or African Green monkeys for which SIV infection is nonpathogenic and does not progress to AIDS despite high-level virus replication^70–72^. Lower frequencies of latently-infected SCM and/or CM CD4+ T cells have also been associated with phenotypes of (i) non-progressing humans maintaining healthy levels of peripheral CD4+ T cells in the absence of ART and (ii) post-treatment HIV controllers, individuals treated soon after infection who controlled viremia after ART discontinuation^73–76^. Altogether, these observations emphasize the importance of preserving long-lived memory CD4+ T cells such as CM CD4+ T cells from HIV infection and persistence.

Despite the transient redistribution of SIV DNA across CD4+ T cell subsets seen during early ART, viral DNA levels were similar in peripheral CD4+ T cells across groups after several months of ART. We acknowledge that the assay used did not distinguish intact from defective proviruses and thus did not measure the true reservoir size. Nevertheless, a reduction of the viral reservoir size through antiproliferative and/or pro-differentiation therapies may require extended or repeated treatment^26^. Interestingly, the observed higher levels of SIV gag RNA in CD4+ T cells in the experimental group as compared to controls suggests Wnt and Notch inhibition promotes a more active reservoir. This result is in line with the increased expression of genes involved in HIV/SIV transcription we observed in CM CD4+ T cells following PRI-724 + LY3039478 treatment.

There were limitations to this study. First, the sample size was limited with 5 to 7 macaques per group. Second, cell sorting was performed using cryopreserved PBMC and EM T cells are particularly vulnerable to freeze-thaw damages. Consequently, small numbers of EM CD4+ T cells were recovered by cell sorting, preventing SIV DNA quantification in some occurrences and thus limiting statistical power. Third, we did not obtain lymph nodes or rectal tissues at matching timepoints between controls and experimental animals and lymphoid tissues were not included in our investigations. Finally, transcriptomic analyses were limited to the CM population which exhibited the greatest changes in response to the experimental treatment and represented a large proportion of the memory pool. Analyses of all maturational subsets may have further elucidated the impact of stemness inhibition on T cell dynamics.

Our experimental approach inhibiting Wnt and Notch pathways during acute SIV infection thus showed a transient redistribution of SIV DNA within the heterogenous CD4+ T cell population likely through enhanced differentiation of self-renewing CD4+ T cells that represent the core of the viral reservoir. Although short-term enhancement of CD4+ T cell differentiation did not reduce the viral reservoir size, the progression of viral DNA into more mature CD4+ T cells is expected to facilitate viral clearance due to their higher turnover rate^8,27^. Careful evaluation of the timing of repeated dosing of Wnt and Notch inhibition is warranted. Furthermore, it has been shown that differentiation into an effector memory phenotype enhances HIV-1 latency reversal in CD4+ T cells *ex* vivo^77–79^. In keeping with these results, our experimental approach promoted a transcriptomic profile consistent with HIV transcription activation in CM CD4+ T cells and led to increased levels of cell-associated SIV gag RNA in total CD4+ T cells suggesting that inhibiting Wnt/Notch signaling may facilitate latency reversal. Future studies using stemness pathway inhibitors in combination with established latency reversing agents will determine their ability to potentiate latency reversal and ultimately facilitate the clearance of the viral reservoir.

## Methods

### Animals, treatments and infection

Five Indian RM (*Macaca mulatta)* with the exclusion of Mamu B*08 and B*17 positive animals, were enrolled in this study. All animals were housed at the Emory National Primate Research Center (Atlanta, GA) and treated in accordance with Emory University and Emory National Primate Research Center Institutional Animal Care and Use Committee regulations (PROTO201900069). Animal care facilities are accredited by the U.S. Department of Agriculture (USDA) and the Association for Assessment and Accreditation of Laboratory Animal Care (AAALAC) International.

For the dose-finding study, two uninfected RMs received two 8-week treatment cycles of PRI-724 administered subcutaneously (s.c.) daily at 20 mg/kg as previously reported^31^, and LY3039478 administered orally (p.o.) at escalating doses of 1.5 and 2.5 mg/kg three times per week. Treatment cycles were separated by an 8-week washout period.

For the efficacy study, all five RMs were infected intravenously (i.v.) with 1x10^5^ tissue culture infective dose (TCID_50_) of SIVmac239 (*nef* open). SIVmac239 stock was titrated *in vitro* for viral infectivity by standard endpoint titration on CEMx174 cells. The 50% tissue culture infectious dose (TCID_50_) was calculated by the method of Reed and Meunch^80^. All animals were immediately initiated on PRI-724 + LY3039478 for 8 weeks as described above. ART treatment was initiated 8 weeks post infection and maintained for the duration of the study. Cryopreserved PBMC samples from a group of 7 ART-suppressed SIV-infected RMs were used for comparative analyses. This ART-only control group received the same viral challenge (virus stock, dose, route) and ART regimen was initiated at the same time post infection as in the 5 RMs treated with PRI-724 + LY3039478. PRI-724 and LY3039478 were administered simultaneously for a period of 8 weeks before ART initiation. PRI-724 was administered s.c. daily at 20 mg/kg. LY3039478 was administered p.o. at 2.5 mg/kg three times per week. The drug ART regimen consisted of two reverse transcriptase inhibitors, 5.1 mg/ml tenofovir disoproxil fumarate (TDF) and 40 mg/ml emtricitabine (FTC), and 2.5 mg/ml of the integrase inhibitor dolutegravir (DTG). The ART cocktail was administered once daily at 1 ml/kg body weight s.c.

### Sample collection and processing

EDTA-anticoagulated blood samples were collected regularly and used for routine assessment of complete blood count, serum chemistry, and immunostaining. Plasma was separated by centrifugation within 1 hour of phlebotomy. Peripheral blood mononuclear cells (PBMCs) were separated by density gradient centrifugation and isolated cells were cryopreserved until use.

### Immunophenotyping by flow cytometry

Multicolor flow cytometric analysis was performed on whole blood or cell suspension using predetermined optimal concentrations of the following fluorescently conjugated monoclonal antibodies (mAbs): CD3-APC-Cy7 (clone SP34-2), CD8-BV711 (clone RPA-T8), CD4-BV650 (clone OKT4), CD95-BV605 (clone DX2), HLA-DR-PerCP-Cy5.5 (clone G46-6), CCR7-FITC (clone 150503), CD45RA-PECy7 (clone 5H9), CD62L-PE (clone SK11) from BD Biosciences, and CD28-PE-Cy5.5 (clone CD28-2) from Beckman-Coulter. Flow cytometric acquisition and analysis of samples was performed on at least 100,000 events on a BD FACS Symphony A5 flow cytometer driven by the FACS Diva software package (BD Biosciences). Analyses of the acquired data were performed using FlowJo version 10.10.0 software (FlowJo, LLC).

### PRI-724 and LY3039478 quantification by LC-MS/MS

Plasma concentration of C-82 (the active metabolite of PRI-724) and LY3039478 were quantified by LC-MS/MS over a 24-hr period following drug administration during the last week of the first and second treatment cycle (PRI-724: 20 mg/kg s.c.; LY3039478: 1.5 mg/kg and 2.5 mg/kg p.o.). Plasma samples were collected at 0, 0.5, 1, 2.5, 6, 8 and 24 hours post-dose. Briefly, 100 μL of plasma was mixed with 20 μL of internal standard (indinavir 125 nM) and extracted with 500 μL of methanol. The supernatant was collected, air-dried, and reconstituted in 200 μL of 50% methanol prior to LC-MS/MS analysis. Samples were further diluted five-fold for quantification of C-82. Chromatographic separation was performed using a Vanquish Flex UHPLC system couple to a Thermo TSQ Quantiva triple quadrupole mass spectrometer (Thermo Fisher Scientific, Waltham, MA) equipped with an electrospray ionization interface. Analytes were separated on a Kinetex XB-C8 column (50 x 2.1 mm, 2.6 μm; Phenomenex, Torrance, CA) using gradient elution with mobile phase A (0.1% formic acid) and mobile phase B (acetonitrile). The LC gradient started with 10% of mobile phase B for 1 min, which then increased from 10 to 90% in 1.5 min and kept at 90% for 1.5 min before returning to the initial condition. Selected reaction monitoring in positive mode was used to detect C-82 (579.2 → 142) and LY3039478 (465.2 → 270). Data were collected and processed by Thermo Xcalibur 3.0 software.

### Amyloid-β peptide quantification by ELISA

Plasma levels of amyloid-β peptide were measured by Enzyme-linked immunoassay (ELISA) over a 24-hour period following drug administration during the first and last week of the second treatment cycle as previously described^81^. Plasma samples were collected at 0 minute, 30 minutes, 1, 2.5, 6, 8 and 24 hours. Plasma amyloid β (1-40) peptide concentration was measured according to the manufacturer’s instruction (IBL-America LTD). All samples were measured in duplicate on a 50 TS Microplate Reader (BioTek, Winooski, VT) at 450 nm. The concentration was calculated based on standard curve, and the mean was taken as sample concentration.

### Plasma SIV RNA quantification

Standard SIVmac239 plasma viral load quantification was performed regularly throughout the study in the Virology Core of the Emory Center for AIDS Research using a standard quantitative real-time PCR (qPCR) assay (limit of detection: 60 copies/ml plasma) as described previously^82^.

### Cell sorting

Prior to FACS sorting, peripheral CD4+ T cells were negatively selected from frozen cell suspension using magnetically labeled microbeads and subsequent column purification according to the manufacturer’s protocol (Miltenyi Biotec). Enriched peripheral CD4+ T cells were then stained with previously determined volumes of the following fluorescently conjugated mAbs: CD3-AF700 (clone SP34-2), CCR7-FITC (clone 150503), CD4-BV650 (clone OKT4), CD8- APC-Cy7 (clone SK1), CD45RA-PE-Cy7 (clone 5H9), CD95-PE-Cy5 (clone DX2) from BD Bioscience; CD28-ECD (clone CD28.2) from Beckman Coulter. Populations of CD4+ T cells for sorting were defined as naïve: CD45RA+CCR7+CD95-; SCM: CD45RA+CCR7+CD95+CD28+; CM: CD45RA-CD95+CCR7+CD28+; TM: CD45RA- CD95+CCR7-CD28+, and EM: CD95+CCR7−CD28-. Sorting was performed on a FACSAria LSR II (BD Biosciences) equipped with FACS Diva software. The sorted cells were flash frozen as a dry pellet and stored at -80°C until further use.

### Cell-associated SIV DNA/RNA quantification in total CD4+ T cells

Cell-associated SIV RNA and DNA was measured simultaneously in peripheral CD4+ T cells (8x10^5^ – 1 x10^6^ cells) lysed in buffer RLT plus (Qiagen) with 2-mercaptoethanol. Nucleic acids were extracted using the Allprep DNA/RNA minikit (Qiagen) according to the manufacturer recommendations with an on-column DNase digestion step. Cell-associated SIVmac239 gag DNA was quantified by qPCR using a 5’ nuclease (TaqMan) assay with SIV *gag* primers, and results were normalized to the RM albumin gene, as described previously^83^. For the quantification of cell-associated RNA, RNA was reverse transcribed using the High-Capacity cDNA Reverse Transcription kit (Thermo Scientific) and random hexamers. SIV gag and the RM CD4 gene were quantified by qPCR of the resultant cDNA using TaqMan Universal Master Mix II (Thermo Scientific). The CD4 forward primer was 5’ -ACA TCG TGG TGC TAG CTT TCC AGA-3’, reverse primer was 5’-AAG TGT AAA GGC GAG TGG GAA GGA-3’, and probe was 5’-AGG CCT CCA GCA CAG TCT ATA AGA-AAG-AGG-3’. The means from two replicate wells were used in all analyses.

### Cell-associated SIV DNA quantification in sorted CD4+ T cells subsets

Frozen sorted CD4+ T cells were lysed with proteinase K (200 µg/ml in 10 mM Tris-HCl [pH8]) for 4 h at 56°C. Simultaneous quantification of SIVmac239 gag DNA and 2-LTR was performed by quantitative multiplex PCR using the 5’ nuclease (TaqMan) assay with an ABI7500 system (PerkinElmer Life Sciences). The sequence of the forward primer for SIVmac239 gag was 5’-GGT TGC ACC CCC TAT GAC AT -3’, the reverse primer sequence was 5’- TGC ATA GCC GCT TGA TGG T -3’, and the probe sequence was 5’-6-carboxyfluorescein-(FAM) AAT CAG ATG TTA AAT TGT GTG GGA-ZEN- 3’ IowaBlack FQ. The sequence of the forward primer for SIVmac239 2-LTR was 5’- CGC CTG GTC AAC TCG GTA CTC-3’, the reverse primer sequence was 5’- GGT ATG ATG CCT TCT TCC TTT TCT AAG-3’, and the probe sequence was 5’-5 2′-chloro-7′phenyl-1,4-dichloro-6-carboxy-fluorescein (SUN) CCC TGG TCT GTT AGG ACC CTT TCT GCT TTG- ZEN- 3’ IowaBlack FQ. For cell number quantification, the quantitative PCR target was the rhesus RPP30 gene which encodes a highly conserved sequence for a ribonuclease enzyme in human and nonhuman primates^84^. The sequence of the forward primer for RPP30 was 5’- AGG ATG CTC CGG GAG TAT GTA -3’, the reverse primer sequence was 5’- CCT GCT TGT CAC CTA TAT AAC AT -3’, and the probe sequence was 5’-6-carboxyfluorescein-(FAM) TCA AGC TGG GAG ACG GAA GAG TCA GT-ZEN- 3’ IowaBlack FQ. Cell lysate was mixed in 50 µl reaction mixtures containing 1X Platinum buffer, 3.5mM MgCl2, 0.2mM deoxynucleotides triphosphate (dNTP), 200 nM primers, 150 nM, probe, and 2 U Platinum Taq. The reactions were performed on a 7500 fast real-time PCR system (Applied Biosystems) with the following thermal program: 2 min at 95°C, followed by 45 cycles of denaturation at 95°C for 10 s and annealing at 60°C for 30 s.

### Transcriptomics analyses

RNA-Seq analyses were conducted at the Emory National Primate Center Nonhuman Primate Genomics Core Laboratory (https://enprc.emory.edu/nhp_genomics_core/). RNA was purified using Qiagen Micro RNeasy columns, and RNA quality was assessed using an Agilent Bioanalyzer instrument. Total RNA (10 ng) was used as input for mRNA amplification using 5’ template-switch PCR with the Clontech SMART-Seq v4 Ultra Low Input RNA kit according to the manufacturer’s instructions. Amplified mRNA was fragmented and appended with dual-indexed barcodes using Illumina Nextera XT DNA library prep kits. Libraries were validated by capillary electrophoresis on an Agilent 4200 TapeStation, pooled, and sequenced on an Illumina HiSeq 3000 sequencer using 151 bases (single end) at an average read depth of 23 million reads. RNA-Seq reads were aligned to the MacaM version 7.8 assembly of the Indian RM genome (available at https://www.unmc.edu/rhesusgenechip/index.htm). Alignment was performed with STAR version 2.5.2b (https://github.com/alexdobin/STAR), and transcript abundance was estimated using htseq-count v0.6.1p1 (http://htseq.readthedocs.io/). Transcript abundance was estimated during the alignment using the method of htseq-count (http://htseq.readthedocs.io/) Read counts were normalized and differential expression analysis was performed with DESeq2 (https://bioconductor.org/packages/release/bioc/html/DESeq2.html). GSEA was performed using the desktop module available from the Broad Institute (https://www.broadinstitute.org/gsea/).

### Statistical analyses

Graphs and statistical analyses were performed using R (version 4.5.3: R Core Team 2026) or GraphPad Prism software version 11.0.1. Comparisons across groups were performed using two-tailed and multiple unpaired Mann-Whitney U tests. Correlations were determined using the nonparametric Spearman correlation coefficient (package metan v 1.19.0). For all statistical analyses, significance was attributed at P values of less than 0.05. Statistical and differential analysis of RNAseq data were performed with DESeq2.

## Acknowledgements

We thank all members of the Mavigner laboratory for their contributions to this study. We also thank the Emory National Primate Research Center Animal and Research Resource teams, the Children’s Healthcare of Atlanta and Emory University Pediatric Flow Cytometry Core, and the Translational Virology and Reservoir Units of the Virology and Molecular Biomarkers Core of the Emory Center for AIDS Research (CFAR) for their support. Antiretroviral therapy was kindly provided by Gilead and ViiV. Figures were created with BioRender. The authors of this paper remember with esteem their colleague and co-author, Dr. S.J. Hurwitz, who passed away on November 23, 2024.

## Authors contributions

MM conceptualize the studies; RRH and IR contributed equally to this work; JN, AL, and IR collected and processed macaque samples; RRH, IR, NS and ACo acquired and analyzed data; ST and SJH performed PK analyses; GKT and SEB generated and analyzed transcriptomic data; RRH, IR and MM interpreted the data and wrote the manuscript; MM and ACh supervised the project; RRH, IR, MM, ACh, SEB, RFS, and GS reviewed and edited the manuscript.

