## Supplementary figures for "Inhibition of Stemness Pathways during Acute SIV Limits Infection of Central Memory CD4+ T Cells and Alters Viral Reservoir Activity in Macaques"

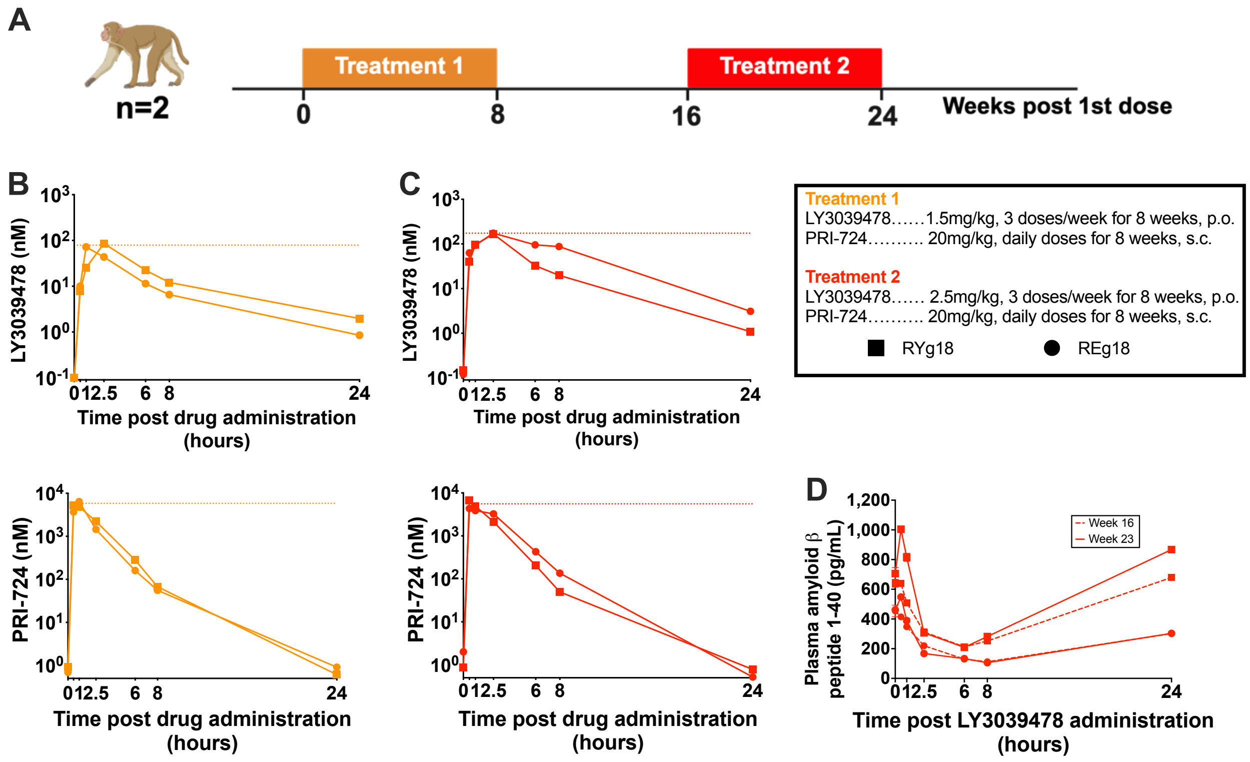


**Supplementary Fig 1. Pharmacokinetic and pharmacodynamic profiles of PRI-724 + LY3039478 administration in uninfected RMs.** (**A**) Experimental design. Two uninfected RMs received 8-week treatment cycles consisting of daily subcutaneous (s.c.) injections of PRI-724 at 20 mg/kg and oral (p.o.) administration of LY3039478 at escalating doses of 1.5 and 2.5 mg/kg three times a week with an 8-week washout period. **B-C** Plasma levels of C-82 (active metabolite of PRI-724) and LY3039478 were measured by LC-MS/MS over 24 hours following administration of PRI-724+LY3039478 during the last week of the (**B**) first or (**C**) second treatment cycle. Horizontal dotted lines represent the average $C_{max}$ for both RMs. (**D**) Plasma level of the amyloid-β peptide measured by ELISA 24 hours following the first dose administered at week 16 and at week 23 of LY3039478 administration during the second treatment period.

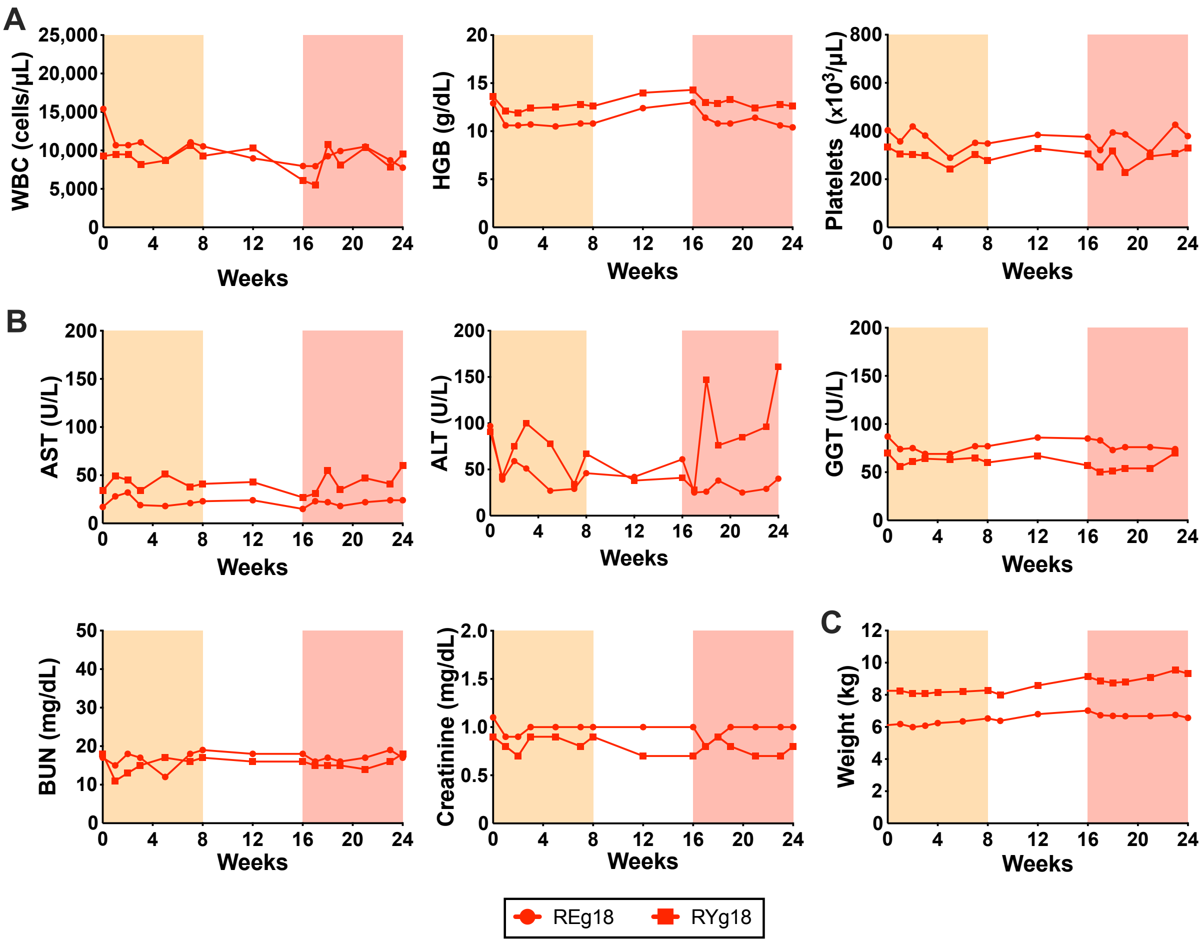


**Supplementary Fig 2. Safety profile of PRI-724 + LY3039478 combined treatment in uninfected RMs. A-C** Longitudinal assessment of (**A**) complete blood counts, (**B**) serum chemistries, and (**C**) weight. WBC: White blood cells; HGB: Hemoglobin; AST: Aspartate aminotransferase; ALT: Alanine transaminase; GGT: Gamma-glutamyl transferase; BUN: Blood urea nitrogen. Shaded areas represent combined treatment cycles (orange: LY3039478 at 1.5 mg/kg and red: LY3039478 at 2.5 mg/kg).

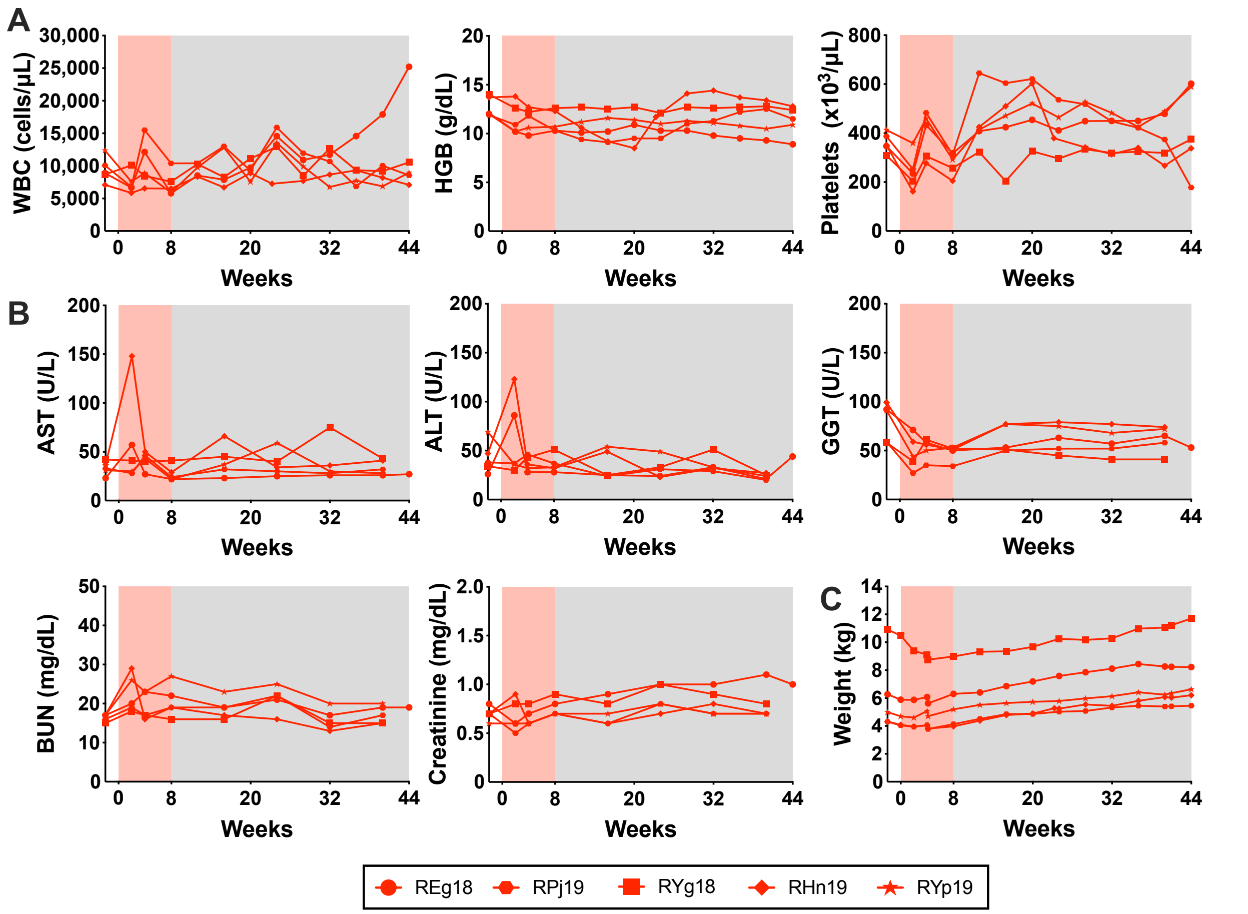


**Supplementary Fig 3. Safety profile of PRI-724 + LY3039478 combined treatment in SIV infected RMs. A-C** Longitudinal assessment of (**A**) complete blood counts, (**B**) serum chemistries and (**C**) weight. WBC: White blood cells; HGB: Hemoglobin; AST: Aspartate aminotransferase; ALT: Alanine transaminase; GGT: Gamma-glutamyl transferase; BUN: Blood urea nitrogen. The red shaded area represents the combined treatment PRI-724 + LY3039478. The grey shaded area represents the period of ART administration.

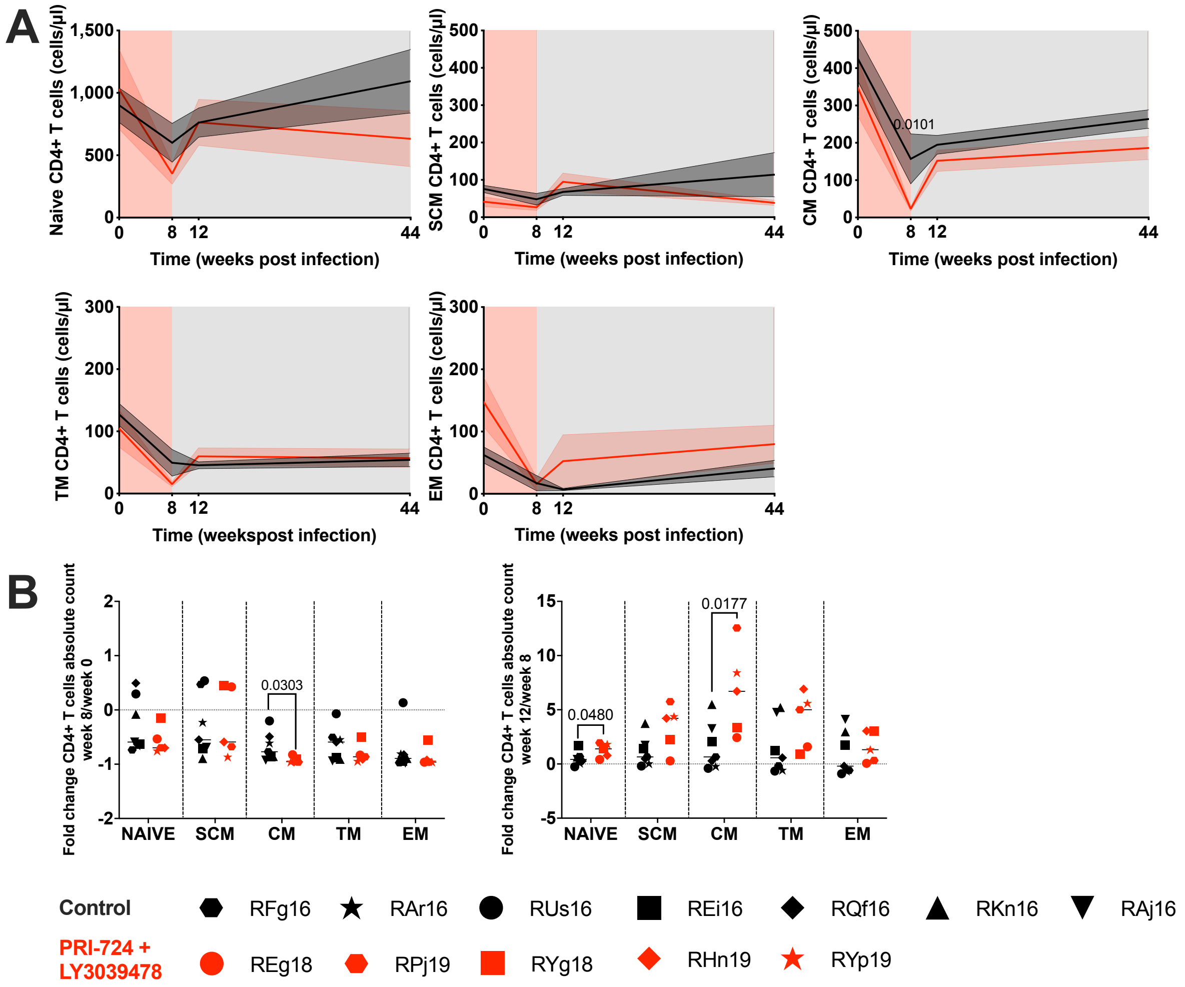


**Supplementary Fig 4. Impact of PRI-724 + LY3039478 combined treatment on CD4+ T cell subset absolute count.** (**A**) Longitudinal assessment of the absolute number of each peripheral CD4+ T cell subset in PRI-724 + LY3039478-treated RMs (red) versus controls (black). Data is represented as mean ± standard error of the mean. (**B**) Fold change in CD4+ T cell absolute counts between 0 and 8, and between 12 and 8 week post infection. Horizontal lines represent the median. A two-sided Mann-Whitney U-test was used to compare values between groups.

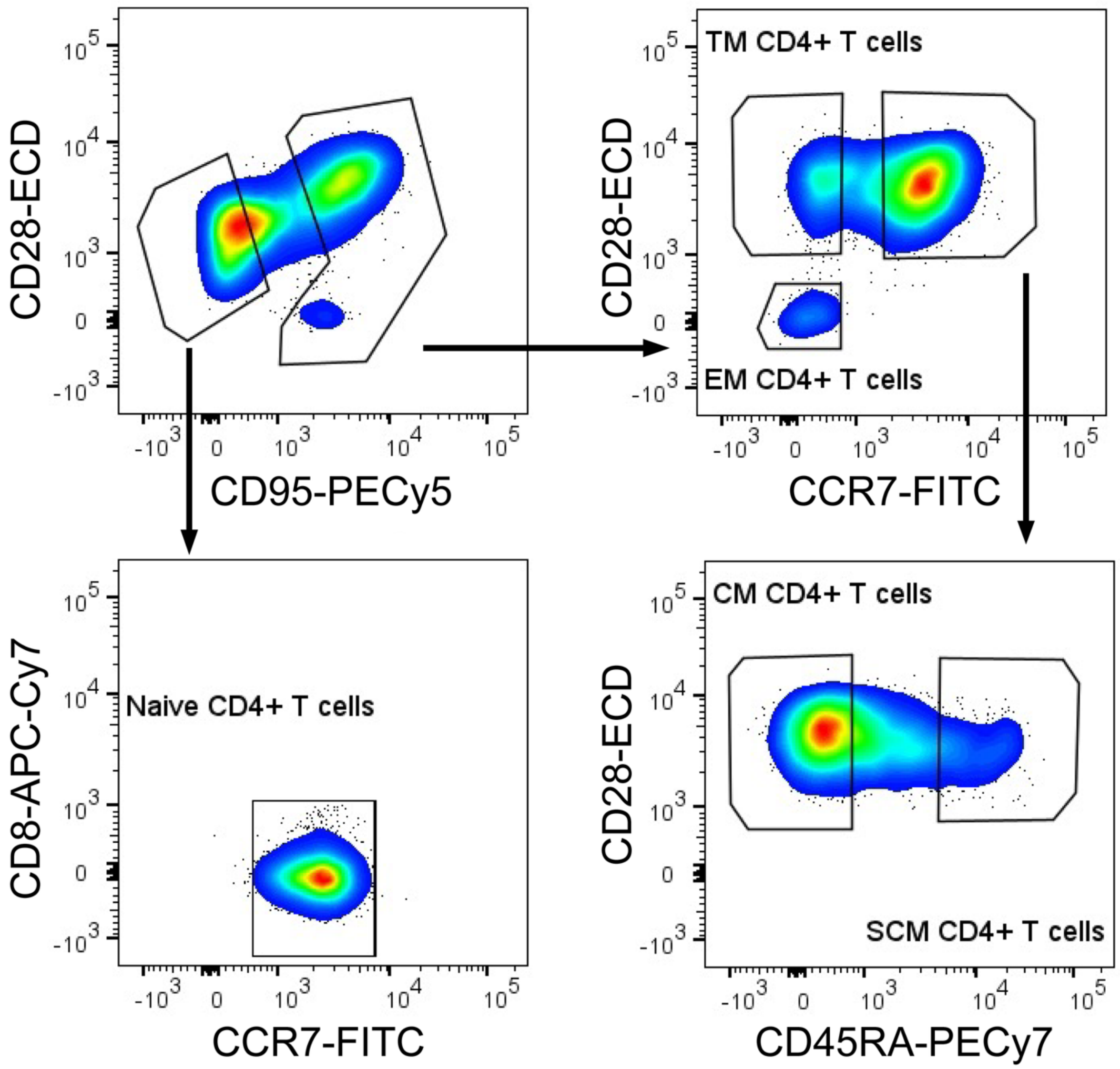


**Supplementary Fig 5. Gating strategy for cell sorting of CD4+ T cell subsets.** Representative flow cytometry dot plots showing the gating strategy used for cell sorting of circulating naïve (CD95-CCR7+), stem cell memory (SCM: CD95+CCR7+CD45RA+), central memory (CM: CD95+CCR7+CD45RA-), transitional memory (TM: CD95+ CCR7- CD28+) and effector memory (EM: CD95+CCR7-CD28-) CD4+ T cells.

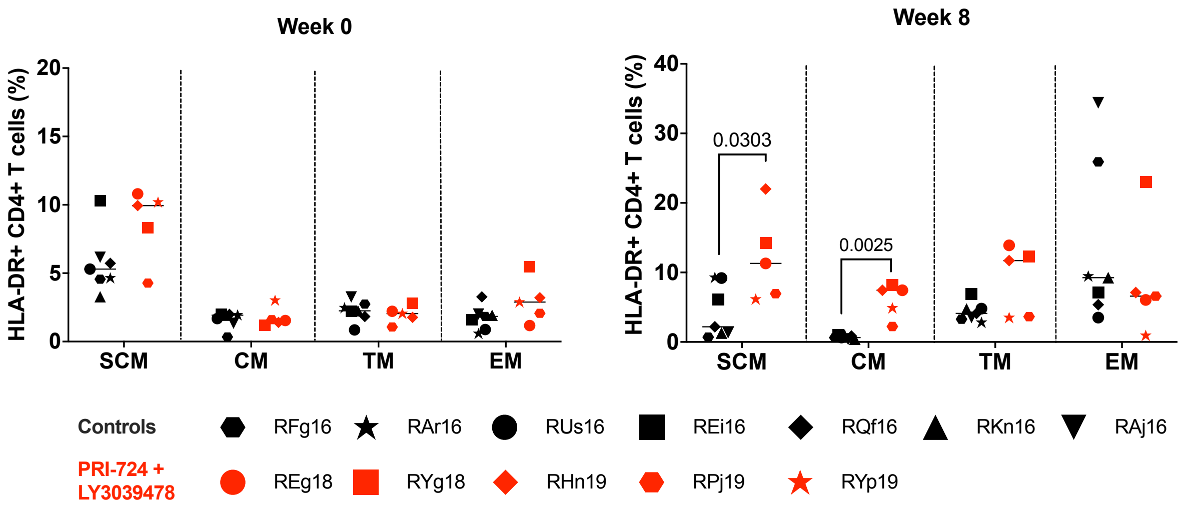


**Supplementary Fig 6. Activation status of memory CD4+ T cell subsets over the stemness inhibition treatment period.** The frequency of memory CD4+ T cells expressing HLA-DR is shown at 0 and 8 weeks post infection. Horizontal bars represent the median. A two-sided Mann-Whitney U-test was used to compare values between groups.

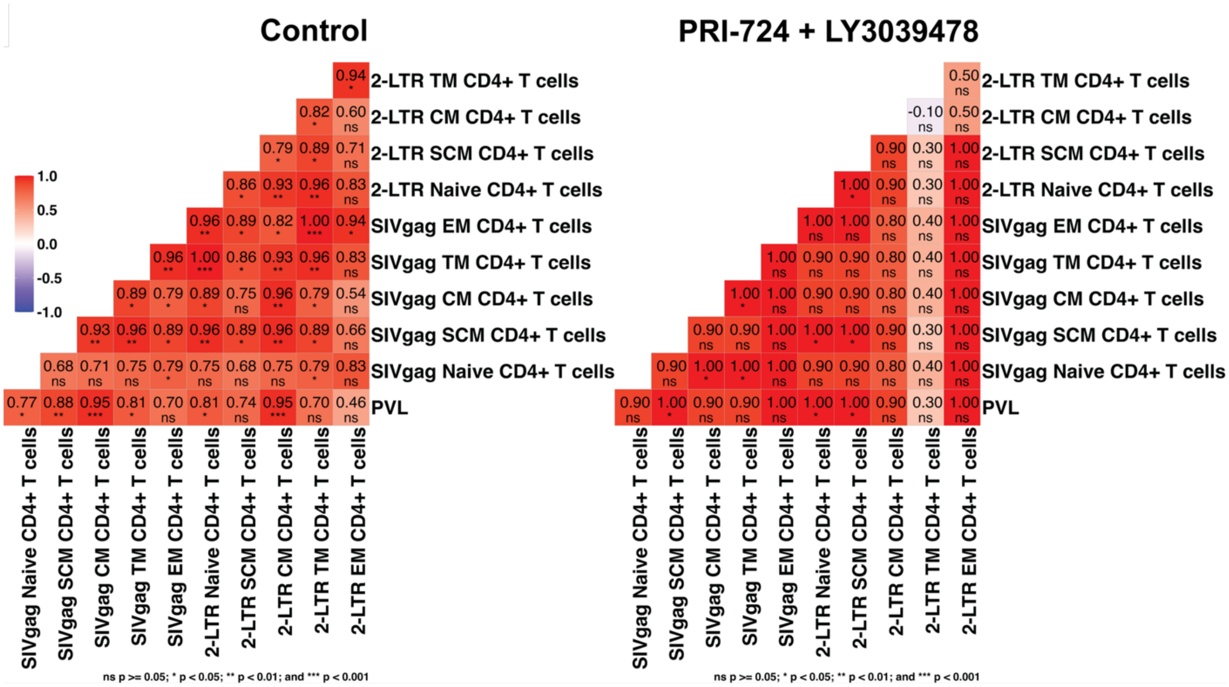


**Supplementary Fig 7. Correlations between virological parameters during untreated SIV infection.** Correlation heatmaps in the control (left) and PRI-724 + LY3039478-treated group (right) at 8 weeks post infection. A non-parametric two-tail spearman correlation was used to generate analyses.

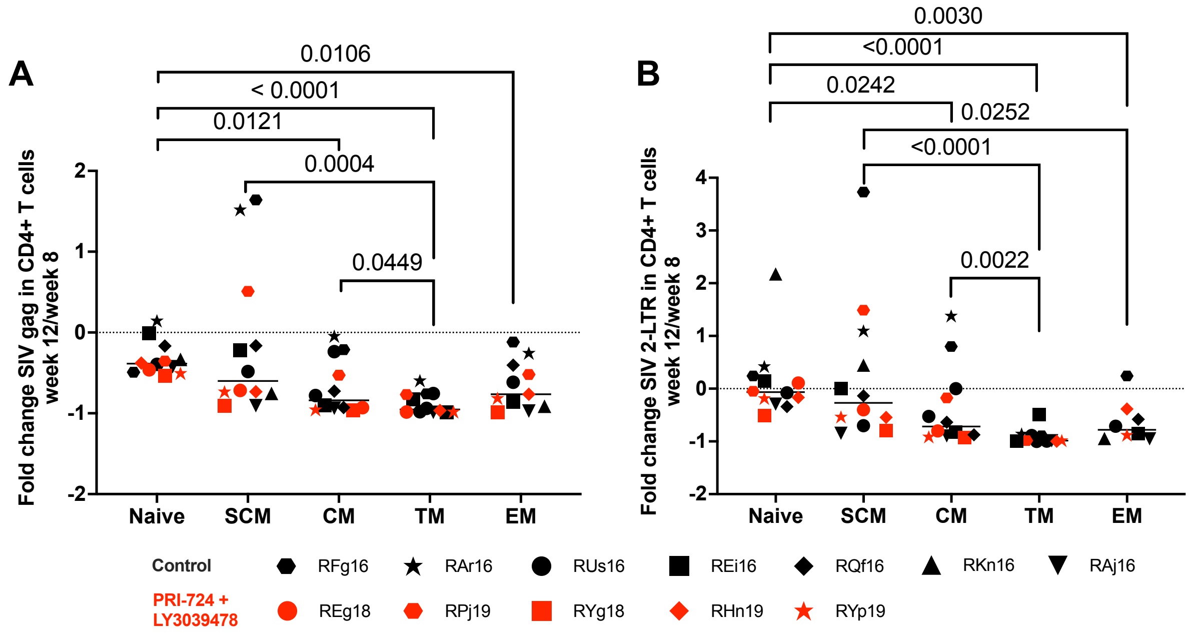


**Supplementary Fig 8. Impact of stemness inhibition on the total and unintegrated SIV DNA after ART initiation.** Fold change in SIV (**A**) gag and (**B**) 2-LTR DNA levels between 8 and 12 weeks post infection in both group in sorted subsets of peripheral CD4+ T cells. A two-sided Mann Whitney U-test was used to compare values between groups.
